# Maternal type 2 immunity promotes a microchimeric transfer of systemic type 2 immunity to offspring

**DOI:** 10.1101/2025.01.03.631175

**Authors:** Malika Gabier, Matthew Darby, Alisha Chetty, Aisha Taliep, Peter Cook, Bernhard Ryffel, Benjamin Dewals, Menno Oudhoff, Adam Cunningham, Emer Hickey, William Horsnell

## Abstract

During maternity mothers undergo an immune pivot to a type 2 immune phenotype which is independent of any antigen experience. In this study we present this maternal Type 2 immunity as a key enabler of optimal maternally-driven microchimeric transfer of immunity to helminth infection in offspring.

To demonstrate that maternal type 2 immunity dictates offspring immunity we nursed wildtype (WT) offspring on WT or IL-4Rα^-/-^ foster mothers. WT offspring nursed on IL-4Ra^-/-^ mothers acquired a reduced type 2 immune signature compared to WT offspring nursed on WT mothers. This demonstrated maternal IL-4Rα imprints type 2 immunity in offspring. This increased type 2 immunity in offspring related to a maternal IL-4Rα dependent increased frequency of maternal microchimeric cells (MMc) being detected in offspring. Higher worm burdens were detected in offspring nursed on IL-4Rα^-/-^ mothers, demonstrating that an antigen independent promotion of maternal type 2 immunity provides offspring with protective immunity against a helminth infection. To establish the contribution of increased MMc in offspring nursed on WT mothers in the control of infection, we undertook an antibody mediated depletion of MMc cells in WT offspring. This MMc depletion impaired the control of infection and reduced the magnitude of offspring type 2 immune response against a helminth infection. These findings present antigen independent maternal IL-4Rα driven type 2 immunity during pregnancy as critical for imparting a profound immune influence via MMc on offspring immunity to infection.

## Introduction

Maternal transfer of immunity provides offspring with high levels of protection against infections in early life^1^. This raised protection is often considered to primarily result from maternal transfer of pathogen specific antibody *in utero* or via nursing^2–4^. However, other immune system components, such as lymphocytes, are also transferred. The importance of the transfer of maternal cells is not fully understood but has been shown to exert profound influence offspring immunity.

We have previously shown that when mothers experience a Type 2 polarising infection with the helminth *Nippostrongylus brasiliensis*, their offspring acquire high levels of pathogen specific protective immune transfer from their mother. This protection was long lasting, antibody independent and associated with offspring acquiring maternal Type 2 CD4+ T cells^5^.

Transfer and tolerance of maternal cells into offspring is called maternal microchimerism (MMc) ^6,7^. MMc can be a powerful driver of offspring immunity and can promote effective transfer of protection from infection to offspring, this can be either dependent^1^ or independent^8^ of a mothers prior antigenic/infectious history. This strong, yet complex, effect of MMc on offspring immunity occurs in spite of the low numbers of cells transferred from the mother and subsequently tolerated by her offspring^7^.

What maternal factors regulates the quality and quantity of MMc to offspring is not well understood. A potentially important regulator of MMc may be the changes that occur to a mothers immunity during pregnancy. One clear change that occurs to maternal immunity during pregnancy is that mothers acquire a Type 2 immune bias^9,10^. This antigen independent maternal type 2 immunity appears to be a beneficial effect of maternity, protecting against important adverse outcomes such as pre-eclampsia^11^. If antigen independent maternal Type 2 immunity alters quantity and quality of MMc derived by offspring is unknown.

In the study presented here, we demonstrate that maternal type 2 immunity influences the control of infection in offspring independently of maternal infection. We present findings showing a profound antigen independent, but maternal type 2 dependent, promotion of offspring immunity that associates with MMc. We then demonstrate this MMc as a key driver of a systemic type 2 immune signature in offspring that contributes to high levels of immunity against early life *N. brasiliensis* helminth infection

## Results

### Pregnancy induces a Type 2 immune profile in WT mothers, but this is reduced in IL-4Rα^-/-^ nursing dams

We first established if pregnancy and type 2 immune competency influenced immunity in female mice. Raised maternal Type 2 immunity during pregnancy was demonstrated by comparing immune readouts between 12-week-old virgin and nursing mothers (14 days post-partum) (Figure 1A). Here the detection of IL-13+ CD4+ T cells in the axial lymph nodes (AxLN) was significantly increased in nursing mothers (Figure 1B) when compared to virgin mice. Additionally, raised numbers of Type-2 innate lymphoid cells (ILC2) were found in nursing mothers in comparison to virgin females (Figure 1C).

**Figure 1.**
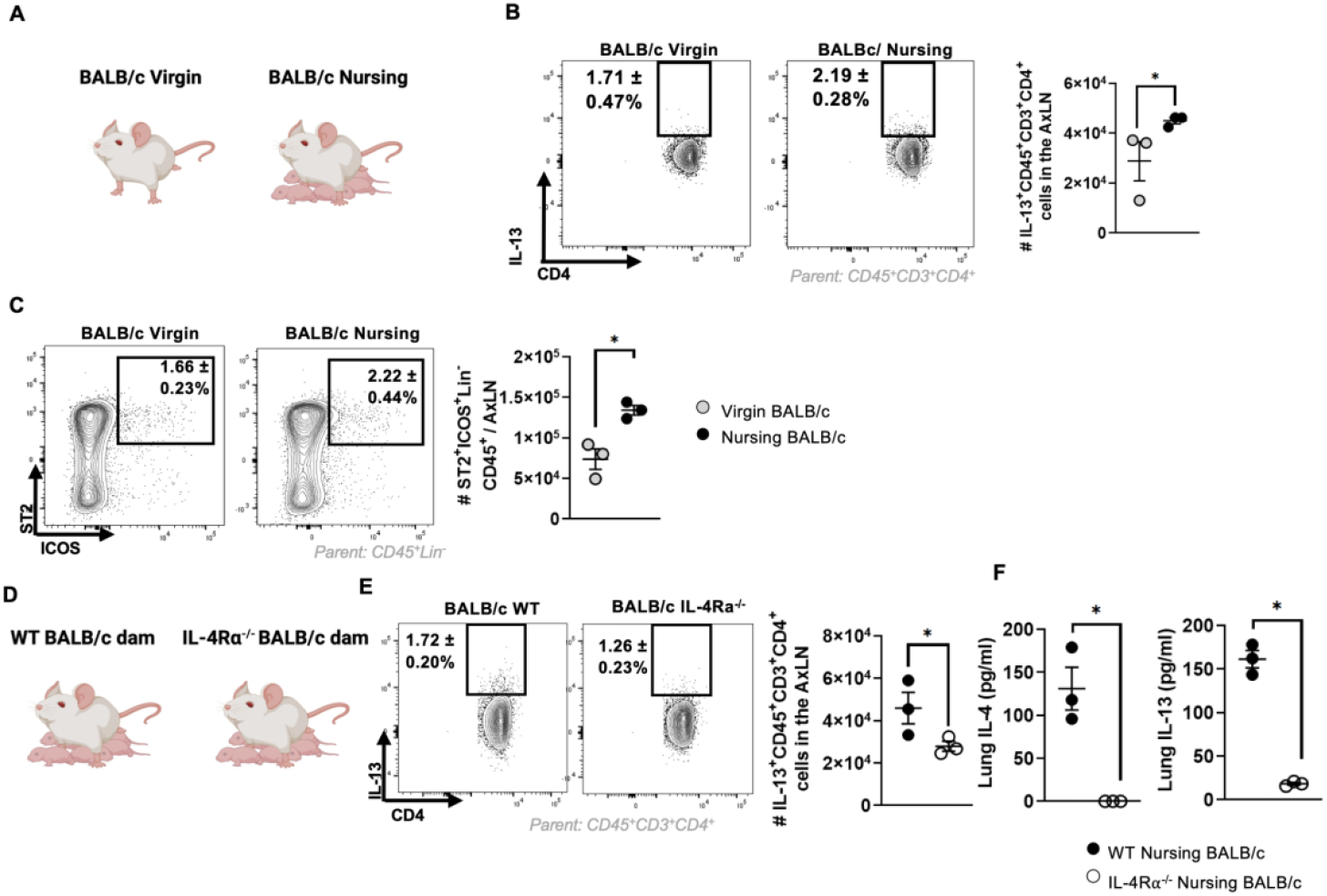
Pregnancy induces a Type 2 immune profile in WT mothers, but this is reduced in IL-4Rα^-/-^ nursing dams. **A** Schematic of virgin vs nursing experiment. BALB/c females were mated and proceeded to nurse their pups. Two weeks after giving birth they were euthanized. Immune readouts were compared to virgin, age-matched female mice. **B - C** AxLN immunity was compared between the nursing and virgin mice. (B) AxLN CD4 T cells expressing IL-13 (IL-13^+^CD45^+^CD3^+^CD4^+^) after stimulation with PMA for 4 hours. (C) Total ILC2 cells per AxLN (CD45^+^Lin^-^ST2^+^ICOS^+^). **D** Schematic comparing WT and IL-4Rα BALB/c nursing foster WT offspring. WT and α IL-4R ^-/-^ females were mated and gave birth to offspring, which were then litter swapped with WT offspring of a similar age 3 days after birth. Foster offspring were nursed until they reached 14-days old, hereafter, mothers and foster offspring were euthanized. **E-F** Immunity was compared from 2-weeks post-partum WT/IL-4Rα ^-/-^ nursing dams. (E) AxLN CD4+ T cells expressing IL-13 (IL-13^+^CD45^+^CD3^+^CD4^+^) after stimulation with PMA for 4 hours. (F) Lung tissue homogenate levels of IL-4 and IL-13. All data are representative of three experimental repeats. Data information: Data are representative of at least two experimental repeats. n= 3 mice per group per experiment. Statistical significance was calculated by Mann Whitney t Test ^*^p < 0.05, ^**^p < 0.01, and ^***^p < 0.001.

Comparison of immune readouts between nursing WT and IL-4Rα^-/-^ BALB/c mice (Figure 1D) demonstrated an expected reduction in type 2 immune readouts in IL-4Rα^-/-^ mothers when compared to WT mothers. This included reduced detection of IL-13+ CD4+ T cells from restimulated AxLN (Figure 1E) and reduced detection of IL-4 and IL-13 levels in lung homogenates (Figure 1F).

### Foster offspring nursed by Type 2 immune deficient mothers display diminished systemic Type 2 immune responses

We next established if there was an immune consequence in WT offspring fostered on IL-4Rα^-/-^ mothers when compared to WT offspring fostered on WT mothers. WT BALB/c offspring were fostered from P1 on either WT or IL-4Rα^-/-^ BALB/c mothers and offspring immunity analyzed when the pups were 14 days old (Figure 2A).

**Figure 2.**
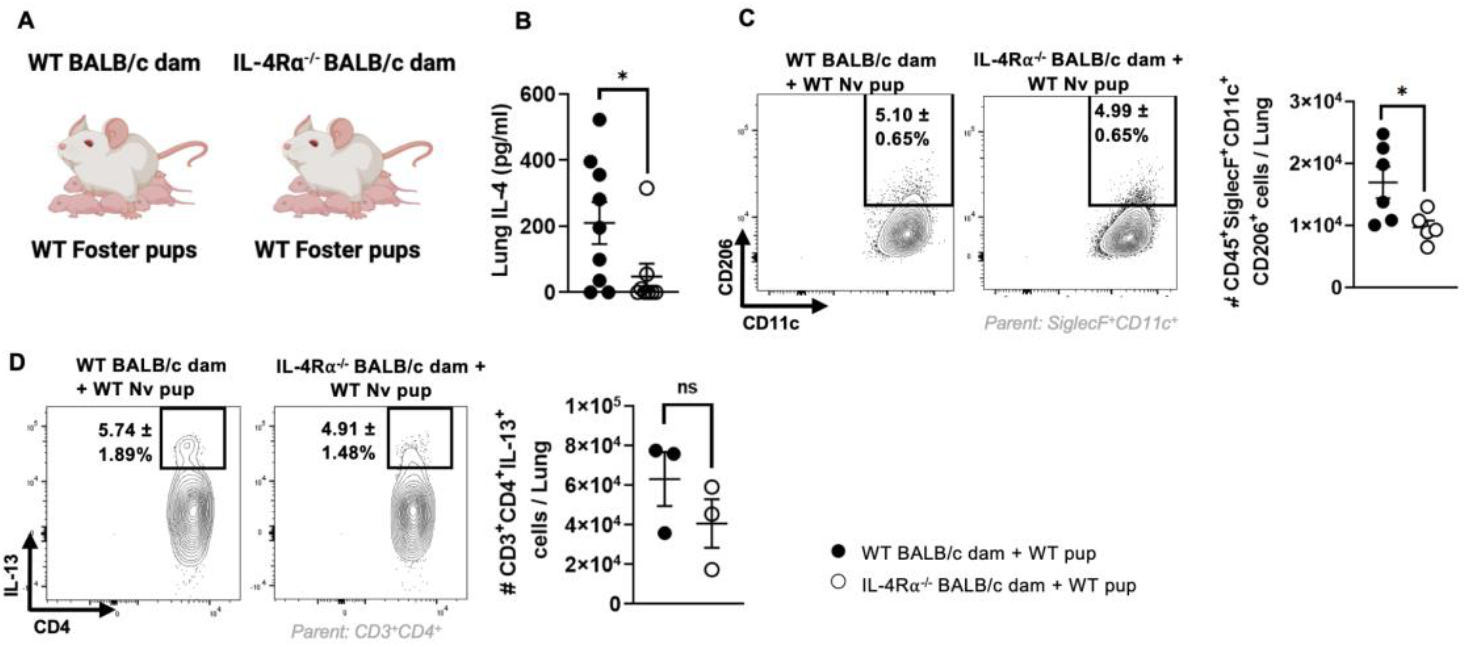
Foster offspring nursed by Type 2 immune deficient mothers display diminished systemic Type 2 immune responses. **A** Schematic of WT/IL-4Rα^-/-^ BALB/c females nursing foster WT offspring. The mice were mated and gave birth to offspring, which were litter swapped with WT offspring of a similar age 3 days after birth. Foster offspring were nursed until they reached 14-days old, hereafter, mothers and foster offspring were euthanized. **B** IL-4 levels in the lung tissue homogenates of 14-day-old WT offspring nursed by WT/IL-4Rα^-/-^ mothers. **C** Total lung M2 alveolar macrophages (CD45^+^SiglecF^+^CD11c^+^CD206^+^) in foster offspring. **D** Total lung CD4+ T cells expressing IL-13 (IL-13^+^CD3^+^CD4^+^) after stimulation with PMA for 4 hours. Data information: Data are representative of at least two experimental repeats. n=3-6 mice per group per experiment. Statistical significance was calculated by Mann Whitney t Test ^*^p < 0.05, ns, not significant.

WT offspring nursed on IL-4Rα^-/-^ BALB/c mothers demonstrated reduced type 2 immune readouts. Here detection of IL-4 in the lung (Figure 2B) and M2 alveolar macrophages (Figure 2C) was reduced in WT offspring nursed on IL-4Rα^-/-^ mothers when compared to WT offspring nursed on WT mothers. However, PMA restimulation of lung cells resulted in no significant reduction in the detection of IL-13+ CD4 T cells in the lung (Figure 2D).

This demonstrates that’s mothers with a profound Type 2 immune deficiency confer a diminished Type 2 immune phenotype to their offspring.

### MMc is reduced in WT offspring nursed on IL-4Rα^-/-^ mothers

Reduced type 2 immunity in pups nursed by WT or IL-4Rα^-/-^ mothers suggests a complex immune influence from the nursing foster mother which may be dependent on maternal transfer of MMc. To address this, we tested if loss of maternal IL-4Rα altered the magnitude of MMc detected in offspring. Here Thy 1.2^-^ BALB/c offspring were fostered and nursed on Thy 1.2^+^ WT or IL-4Rα^-/-^ mothers. Staining for Thy1.2 enables us to identify both maternal-derived (Thy1.2^+^) and endogenous derived (Thy1.2^-^) cells (Fig 3A). Fostered P14 WT pups nursed by IL-4Rα^-/-^ mothers had significantly reduced maternally derived immune cells (CD45+Thy1.2+) in their lungs than pups nursed on WT mothers (Figure 3B). We have previously identified that CD4+ MMc from infected mothers associate with offspring acquiring immunity to infection^5^. In the current study we found that WT offspring nursed on IL-4Rα^-/-^ mothers displayed significantly reduced maternally acquired CD4+ T cells (CD45+CD3+CD4+Thy1.2+) in their lungs (Figure 3C). Therefore, in the absence of maternal IL-4Rα offspring acquire a reduced MMc transfer that is independent of maternal antigen exposure.

**Figure 3.**
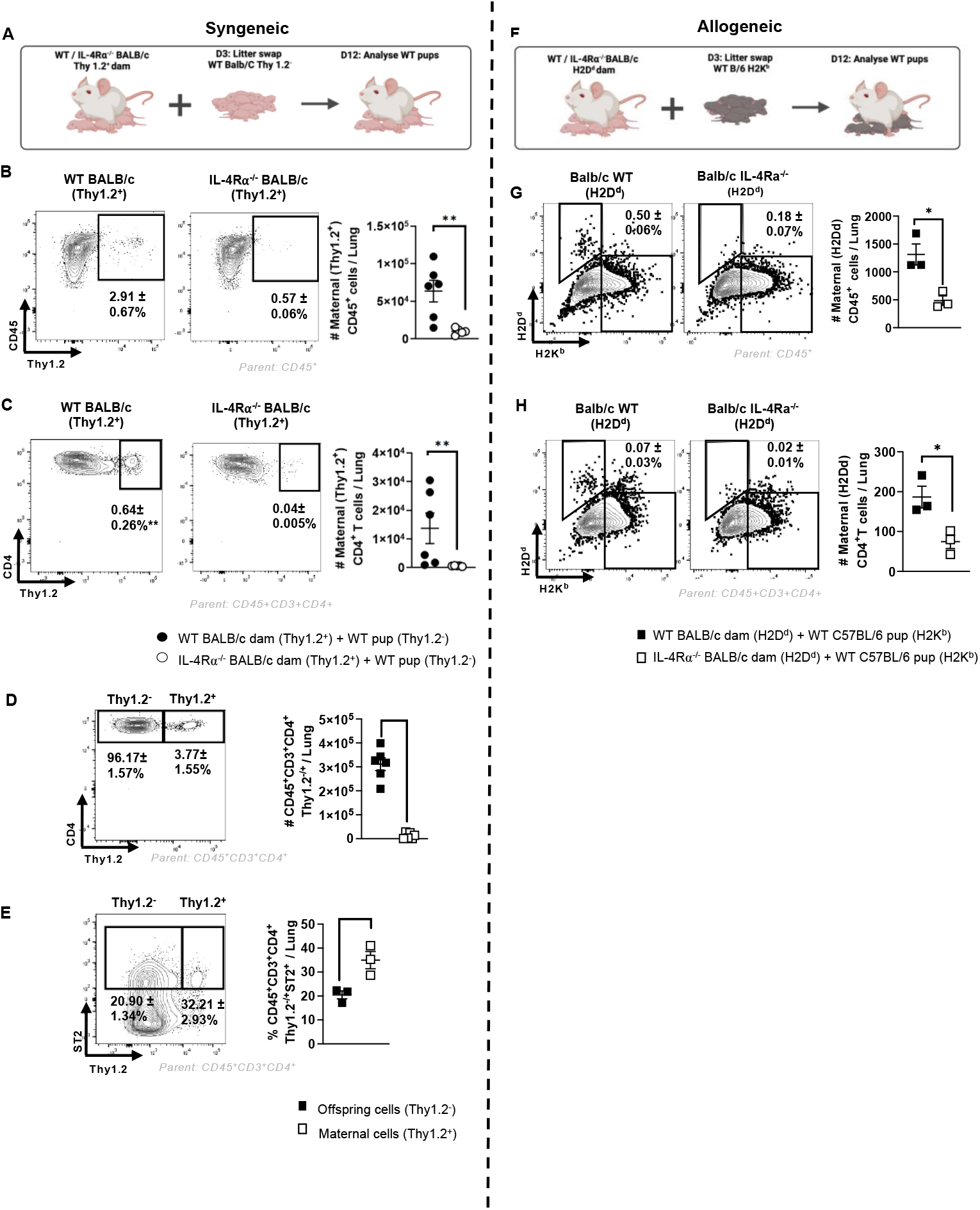
MMc is reduced in WT offspring nursed on IL-4Rα^-/-^ mothers. **A** Schematic of the allogeneic fostering experiment. WT Thy1.2^-^ BALB/c pups were fostered by Thy1.2^+^ WT/IL-4Rα^-/-^ BALB/c mothers at 3 days old. Foster Thy1.2^-^ pups were euthanized at 14 days old and lungs were harvested for immune analysis. **B – E** Analysis of WT foster pups’ lung immunity following nursing on WT or IL-4Rα^-/-^ dams. (B) Measurement of maternally derived CD45+ cells (CD45+Thy1.2+) in the lungs of the foster pups (C) Measurement of maternally derived CD4+ T cells (CD45^+^CD3^+^CD4^+^Thy1.2^+^) in the lungs of the foster pups nursed by WT or IL-4Rα^-/-^ dams. (D) A comparison of endogenous CD4+ T cells (CD45^+^CD3^+^ CD4^+^Thy1.2^-^) and maternally derived CD4+ T cells (CD45^+^CD3^+^CD4^+^Thy1.2^+^) in the lungs of the foster pups nursed by WT or IL-4Rα^-/-^ dams. (E) A comparison of endogenous Type 2 CD4+ T (CD45^+^CD3^+^CD4^+^Thy1.2^-/^ST2^+^) and maternally derived Type 2 CD4+ T cells (CD45^+^CD3^+^CD4^+^Thy1.2^/+^ST2^+^) in the lungs of foster pups. **F** Schematic of the syngeneic fostering experiment. 3-day old WT Bl/6 H2Kb^+^ pups were fostered by WT/IL-4Rα^-/-^ BALB/c H2Dd^+^ mothers. Foster B/6 pups were euthanized at 14 days old and lungs were harvested for immune analysis **G – H** Analysis of WT BI/6 foster pups’ lung immunity following nursing on WT or IL-4Rα^-/-^ BALB/c dams. (G) Measurement of maternally derived CD45+ cells (CD45^+^H2Kb^-^H2Dd^+^) in the lungs of the foster pups nursed by WT or IL-4Rα^-/-^ dams. (H) Measurement of maternally derived CD4+ T cells (CD45^+^CD3^+^CD4^+^H2Kb^-^H2Dd^+^) in lungs of foster pups nursed by WT or IL-4Rα^-/-^ dams. Data information: All data are representative of at least two experimental repeats. n= 3-6 mice per group per experiment. Statistical significance was calculated by Mann Whitney t Test ^*^p < 0.05, ^**^p < 0.01.

To determine how MMc contribute to the overall immune cell population in the pups’ lungs, levels of endogenous and maternally derived CD4+ cells in the lung were compared. This identified that MMc CD4+ T cells contributed to <4% of the lung CD4+ T cell population in this syngeneic foster model (Figure 3D). However, the proportion of maternally derived CD4+ T cells expressing the Type 2 marker ST2 was significantly higher than endogenous ST2+CD4+ T cells (Figure 3E). Thus, maternally derived CD4+ cells have a raised T2 immune bias when compared to offspring CD4+ T cells.

We next tested this in a more physiologically relevant allogeneic setting that replicates mother-offspring MHC mismatch, a critical physiological feature of the mother-offspring dyad. Here WT BL/6 pups (H2K^b^) were fostered by WT or IL-4Rα^-/-^ BALB/c mothers (H2D^d^). In P14 fostered pups the level of MMc in lungs was established (Figure 3F). As in the syngeneic setting, the foster pups nursed by IL-4Rα^-/-^ mothers acquired less maternally derived (H2D^d+^) CD45+ and CD4+ T cells than pups nursed on WT mothers (Figure 3B G and H). Pup groups nursed in an allogeneic setting maintained less maternally derived MHC mismatched immune cells than the pups nursed in the syngeneic setting (Figure 3A).

Thus, nursing mothers with a Type 2 immune deficiency have an impaired ability to transfer MMc to their offspring, demonstrating for the first time, that Type 2 maternal immunity exerts a significant influence on the magnitude of MMc detected in offspring.

### Offspring launch impaired Type 2 immune control of helminth infection if mothers lack IL-4Rα

As WT pups demonstrated reduced type 2 immunity if nursed by mothers lacking IL-4Rα in comparison to WT offspring nursed on WT mothers we sought to test if this had a consequence for control of infection with *N. brasiliensis*. Here 14-day old Bl6 pups fostered on either WT or IL-4Rα^-/-^ BALB/c mothers were infected with *N. brasiliensis* and then analyzed at 8 days post-infection (PI) (Figure 4A).

**Figure 4.**
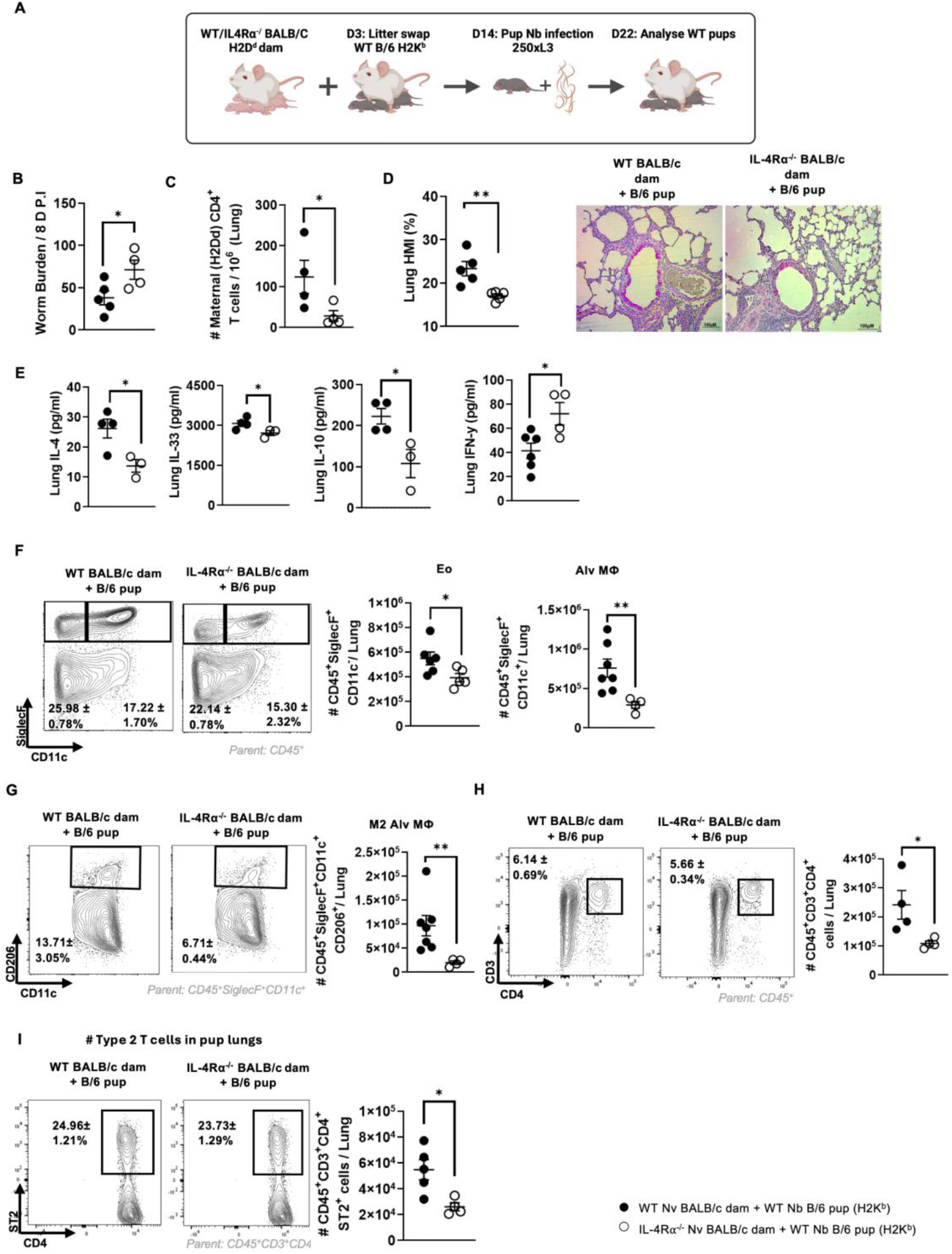
Offspring launch impaired Type 2 immune control of helminth infection if mothers lack IL-4Rα. **A** Schematic of allogeneic fostering experiment with subsequent pup Nb infection. B/6 pups were fostered by naïve WT/IL-4Rα^-/-^ BALB/c mothers at 3 days old. 14-day-old foster pups were subcutaneously infected with 250x L3 *Nippostrongylus brasiliensis* and euthanized 8 days post infection. The pup lungs were harvested for immune analysis. **B** Measurement of maternally derived CD4+ T cells (H2DdCD45^+^CD3^+^CD4^+^) in the lungs of the foster pups. **C** Pup intestinal worm counts at day 8 post infection. **D** Histological mucus index % for quantification of goblet cells in the lung airways of the pups. **E** The levels of IL-4, IL-33, IL-10 and IFN-γ in the pups’ lung homogenates. **F - G** Comparison of eosinophil and alveolar macrophage populations in the lungs of Nb infected pups nursed by WT or IL-4Rα^-/-^ mothers. (F) Measurement of the total eosinophil (CD45^+^SiglecF^+^CD11c^-^) and total alveolar macrophages (CD45^+^SiglecF^+^CD11c^+^) in the pups’ lung. (G) Measurement of the alveolar macrophages expressing CD206 (CD45^+^SiglecF^+^CD11c^+^CD206^+^) in the pups’ lung. **H – I** Comparison of T cell populations in the lungs of Nb infected pups nursed by WT or IL-4Rα^-/-^ mothers. (H) Measurement of total CD4 T cells in the pups’ lung (CD45^+^CD3^+^CD4^+^). (I) Measurement of total Type 2 CD4 T cells (CD45^+^CD3^+^CD4^+^ST2^+^) in the pups’ lung. Data information: Data shown as mean ± SEM from two independent experiments. n = 4-7 mice per group per experiment. Statistical significance was calculated by Mann Whitney t Test *p < 0.05 and **p < 0.01.

WT pups nursed by IL-4Rα^-/-^ mothers demonstrated significantly higher intestinal worm burdens than those seen in WT pups nursed on WT mothers (Figure 4B). Associated with this was reduced detection of MMc CD4+ T cells in offspring nursed on IL-4Rα^-/-^ mothers (Figure 4C).

The offspring nursed by IL-4Rα^-/-^ mothers displayed a reduced ability to launch a protective Type 2 immune response against *N. brasiliensis*. Airway mucus secretion by goblet cells in WT pups nursed by IL-4Rα^-/-^ mothers was reduced when compared to WT pups nursed on WT mothers (Figure 4D). Moreover, the Type 2 (e.g. IL-4 and IL-33) and the regulatory (IL-10) cytokine classically upregulated following *N. brasiliensis* infection were reduced while the Type 1 cytokine IFN-γ was increased in the lungs of WT pups nursed by IL-4Rα^-/-^ mothers when compared to WT pups nursed on WT mothers (Figure 4E).

Additionally, eosinophils and M2 macrophages, Type 2 myeloid cells typically raised following *N. brasiliensis* infection, were reduced in the lungs of WT pups nursed by IL-4Rα^-/-^ mothers when compared to WT pups nursed on WT mothers (Figures 4F and G). This was also the case with CD4+ T cells with both total (CD45+CD3+CD4+) and type 2 CD4 T cells (CD45+CD3+CD4+ST2+) being reduced in WT pups nursed by IL-4Rα^-/-^ mothers when compared to WT pups nursed on WT mothers (Figures 4H and I).

### *In vivo* depletion of MMc in offspring impairs Type 2 responses against helminth infection

Collectively our data demonstrates that maternal Type 2 immunity promotes a Type 2 immune profile in offspring, which is associated with raised detection of MMc lymphocytes in offspring. Moreover, this leads to an enhanced protective immune response against a type 2 resolved infection, namely by the helminth *N. brasiliensis*.

To establish if MMc contributed to raised offspring Type 2 immunity we undertook *in vivo* depletion of maternally derived cells to determine if loss of these cells impaired the pup immune responses against *N. brasiliensis* infection. Here WT Thy 1.2 Bl/6 pups were fostered by WT Thy1.1 BALB/c mothers. At regular intervals, one pup group (referred to as treated) received anti-Thy 1.1 antibody to deplete the MMc *in vivo*. Pups were infected with Nb and after 8 days of infection immune readouts were measured (Figure 5A).

**Figure 5.**
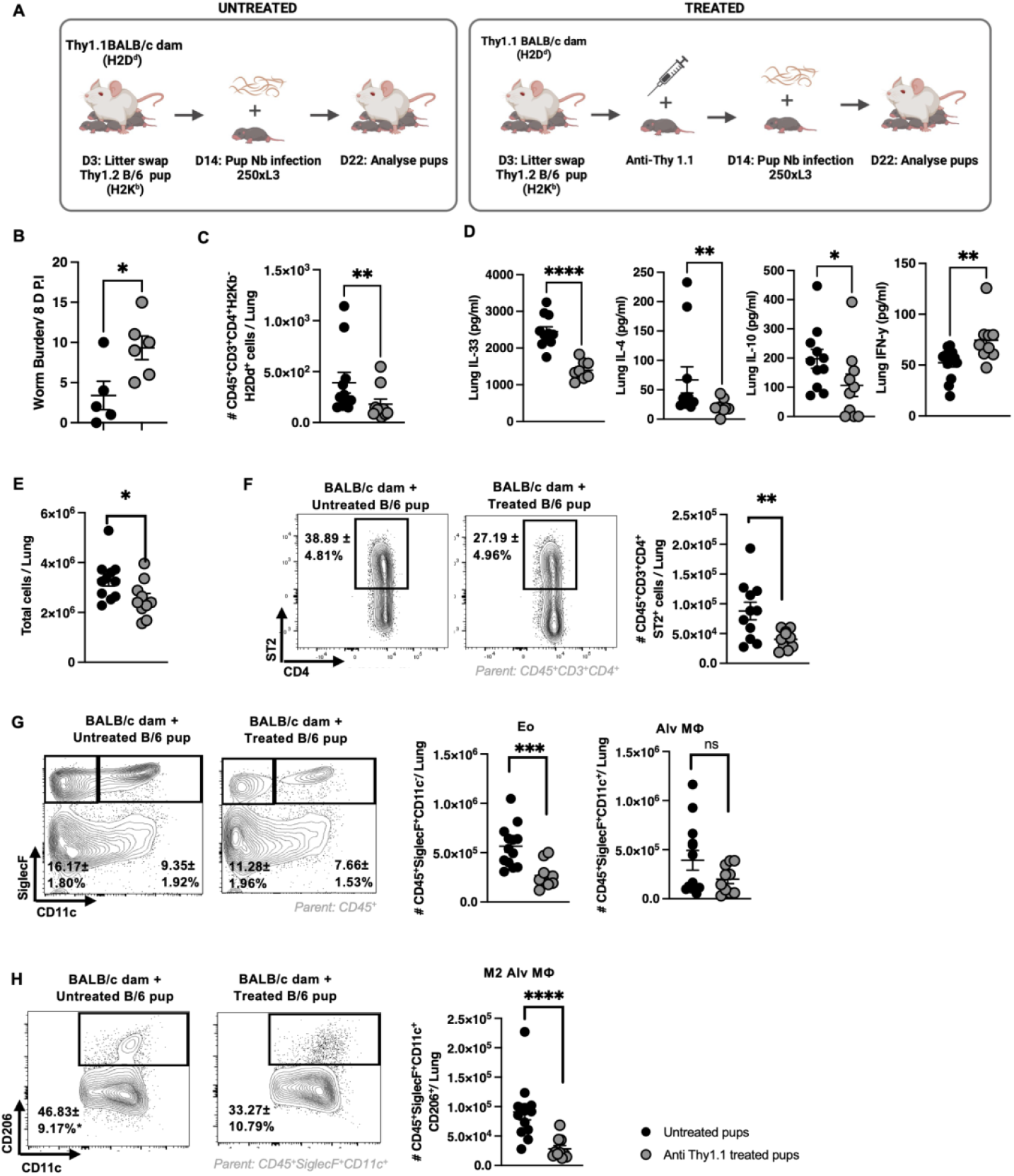
*In vivo* depletion of MMc in offspring impairs Type 2 responses against helminth infection. **A** Schematic of *in vivo* depletion of MMc using anti-Thy1.1. WT Thy1.2 B/6 pups were fostered by WT Thy1.1 BALB/c mothers at 3 days old. On day 13, 15, 17, 19 and 21 ‘treated’ foster pups received 0.25mg Anti-Thy1.1 depletion antibody / PBS to deplete the MMc *in vivo*. On day 14 all pups were subcutaneously infected with 250x L3 *Nippostrongylus brasiliensis* and euthanized 8 days post infection. The lungs were harvested from pups for immune analysis. **B** Pup intestinal worm counts at day 8 post infection. **C** Maternally derived CD4 T cells (CD45^+^CD3^+^CD4^+^H2D^d+^) in the pup’s lungs. **D** The levels of IL-4, IL-33, IL-10 and IFN-γ in the lungs of anti-Thy1.1 treated and untreated pups. **E** Total lung cells per pup. **F** Measurement of total Type 2 CD4 T cells (CD45^+^CD3^+^CD4^+^ST2^+^) in the pups’ lung. **G - H** Myeloid immune cell populations in the lung were compared between the anti-Thy1.1 treated and untreated pups following a Nb infection. (G) Eosinophils and alveolar macrophages in the pups’ lung. (G) M2 alveolar macrophages in the pups’ lung. Data information: Data shown as mean ± SEM from at least two independent experiments. n = 4-6 mice per group per experiment. Statistical significance was calculated by Mann Whitney t Test ^*^p < 0.05, ^**^p < 0.01, and ^***^p < 0.001, ^****^p < 0.0001, ns, not significant

Following infection with *N. brasiliensis* anti-Thy1.1 treated pups demonstrated higher worm burdens (Figure 5B) and displayed a significant reduction in the number of MMc detected in the lung (Figure 5C). This was associated with an impaired ability to launch a Type 2 response to a *N. brasiliensis* infection. Here MMc depleted pups displayed reduced lung IL-4, IL-33 and IL-10 when compared to the untreated mice (Figure 5D). IFN-γ in the lung was elevated in the MMc depleted pups (Figure 5D). Additionally, total number of cells in the lung homogenates of the pups were significantly reduced in MMc depleted pups when compared to untreated pups (Figure 5E). The number of maternally derived CD4+ T cells in pup lungs were also reduced in anti-Thy1.1 treated pups compared to the untreated pups. This was further explored within the context of type 2 immunity which identified the numbers of ST2+ CD4+ cells were reduced in MMc depleted pups (Figure 5F). This effect also extended to myeloid cells with reduced eosinophil and macrophage populations in MMc depleted pups when compared to untreated pups (Figure G and H).

This demonstrates, for the first time, that a competent maternal type 2 immunity enables optimal MMc transfer of type 2 systemic immunity via nursing to offspring. This MMc transferred type 2 immunity provides an antigen independent protection against helminth infection in offspring.

## Discussion

Our study reveals that maternal Type 2 immunity, a prominent immune signature of pregnancy, exerts an antigen independent influence on offspring’s immunity to infection. Moreover, this effect was, to a great extent, exerted by MMc as targeted disruption of MMc resulted in a clear reduction in offspring Type 2 immunity following a helminth infection.

We therefore have demonstrated, for the first time, that maternal Type 2 bias provides a major antigen independent influence on offspring ability to control a helminth infection. This may represent an evolutionary adaptation to protect against this important class of infection in early life, when risk of infection and resulting pathology are at their highest^12–14^. Reports from others have shown MMC can provide antigen independent early life immunity that is beneficial for early life control of viral infection in both humans and mice^8^. However, to the best of our understanding, how underlying maternal immunity influences offspring ability to control infection has not been demonstrated before.

We present immune cell transfer via breastfeeding as a potentially important source of MMc in offspring. While it is well known that breast milk contains a full profile of immune cells including ILCs^15^, T cells^16,17^ and myeloid cells^18,19^ the extent to which these cells influence offspring immunity is not fully appreciated. In our study we identify that MMc transfer via nursing exerts a significant and systemic influence on offspring immunity to infection. Notably our demonstration of reduced protective Type 2 immunity in offspring following targeted depletion of maternal CD90+ lymphocytes implies MMc T cells provide an antigen independent protection against infection.

Together our findings that an inherent change in mothers immunity during pregnancy (Type 2 immune bias) dictates the quality of immune transfer to offspring, represents a paradigm shift in our fundamental understanding of maternal transfer of immunity. A number of human and animal studies have already shown that changes in maternal immunity due to infection can influence offspring immunity^20,21^. We present mothers being capable of transferring high levels of immune protection to offspring that is independent of maternal infectious history. This discovery provides a new layer of understanding of how mothers immunity during pregnancy critically influences the quality of immune transfer derived by her offspring.

This discovery has important relevance clinically. Widely used drugs, such as duplimiab (anti-IL-4Rα) and benralizumab (anti-IL-5) target type 2 immunity to treat allergic disease^22^. It is not understood if use of these drugs by pregnant women has an influence on their children’s immunity. Additionally, functional polymorphisms in IL-4Rα are common^23^. This includes the Q576R polymorphism, which occurs at frequencies of 5% globally and is recognised as inherently raising Type 2 immunity^24^. The impact of such polymorphisms on offspring immunity *per se* is not currently understood. The findings we report here provide a strong justification for addressing the transgenerational immune influences of such drugs and polymorphisms.

In summary our study is the first of its kind to demonstrate that a pronounced pregnancy induced change to the maternal immune system (raised Type 2 immunity) is essential for optimal transfer of immunity to infection to offspring. Moreover, we demonstrate that a key component of this transfer of immunity is maternal microchimerism. Here the maternal type 2 dependent transfer and tolerance of microchimeric cells by offspring provides a striking level of antigen independent immunity against infection. Our data therefore presents a novel mechanism where mothers immunity can train offspring responses to infection via the transfer of immune cells naïve to that pathogen.

## Materials and Methods

### Mice used

The following BALB/c background mouse strains were used in the study: BALB/c (Thy1.2 also called CD90.2), IL-4Rα^-/-25^ and Thy1.1 (CD90.1). The BALB/c mice used express MHC1 haplotype H2K^d^/H2D^d^. C57BL/6 were used which express MHC1 haplotype H2K^b^/H2D^b^. Mice were bred and housed in specific pathogen–free conditions at the Animal Unit of the University of Cape Town, South Africa. All experimental mice were used between 1–15 weeks of age with appropriate littermate controls of the same generation. Mice were euthanized by lethal halothane inhalation. *Ethics statement:* All studies were carried out in accordance with protocol 018/041 or 023/015 approved by the Faculty of Health Sciences Animal Ethics Committee from the University of Cape Town

### Mating and litter-swaps

Female mice aged between 6 and 7 weeks were mated with male mice at one male per cage of two females for seven days, after which the male was removed. Females gave birth approximately 21 days after fertilisation and birth of pups was monitored daily to ensure age matching. Three days after birth, age-matched pups were gently removed from their birth mother and transferred to a foster mother. Pups were nursed until they reach 14 days old (unless specified differently). A maximum of 2 mothers and 7 pups were accommodated in a single cage.

### Infection with Nb

14 day old pups were infected subcutaneously with 250 x *N. brasiliensis* L3 larvae. To enumerate adult worms, mice were euthanised 8 days post-infection (PI), and intestines opened longitudinally, incubated in 10 ml 0.9% saline for 3 hrs at 37°C. Following incubation, parasites were counted under a dissecting microscope.

### In vivo depletion of MMc with anti-Thy1.1

To deplete MMc cells acquired by offspring via nursing, foster offspring (Thy 1.2) were injected intra-peritoneally with 0.25mg Anti-Thy1.1 or isotype control (Clone: 19E12; BioXcell) every 48h from P12 until end of experiment. CD90 (Thy1.1 or Thy1.2) is found on T cells and ILCs. Thus these two cell populations will be targeted for depletion by the treatment.

### Histology

Tissue samples were fixed in 10% neutral buffered formalin solution, dehydrated and embedded in paraffin according to standard protocols. PAS (Periodic acid-Schiff) staining was performed according to standard protocols on 5µm sections to identify mucous production, with PAS+ Goblet cells presenting with a deep magenta colour Images at 10X magnification were obtained on a Zeiss Primostar 1 light microscope and analysed with ImageJ analysis software.

### Flow Cytometry

Single cell suspensions were prepared, and 1 × 10^6^ cells incubated in PBS + 0.5% BSA, 1% normal rat serum and appropriate antibody cocktails. Cell populations were determined and acquired on a BD FACS Fortessa (Becton Dickinson). Cell populations were identified by the following antibody staining strategies; CD4 T cells: CD3+CD4+. CD4 T cells populations were additionally stratified into activated effector (CD44+CD62L_lo_), type 2 T1/ST2+, and Thy1.1+Thy1.2- or Thy1.2+Thy1.1- or H2K^b+^H2D^d-^ or H2K^b-^H2D^d+^ (maternal vs endemic) T cell populations. ILC2s: CD45+Lineage^-^ CD127+ICOS+. Eosinophils: CD45+SiglecF+CD11c-. M2 alveolar macrophages: CD45+SiglecF+CD11c+CD206+. CD45 cells: CD45+

Intra-cellular cytokine staining was also carried on T cells to determine IL-13 expression. Cells were re-suspended in complete media (IMDM (GIBCO/Invitrogen; Carlsbad, CA) + 10% FCS, P/S) at 2.5×10^7^/ml and stimulated with 10 μg/ml PMA/ionomycin and GolgiStop (as per manufacturer’s protocol; BD Pharmingen) at 37°C for 4 hours. After re-stimulation, cells were surface stained for CD3 and CD4 then fixed and permeabilised with Cytofix/Cytoperm Plus (as per manufacturer’s instructions; BD Pharmingen). Intracellular staining was performed by staining cells with IL-13 or appropriate control. All analyses were performed with FlowJo© software.

### Cytokine ELISA

Cytokine ELISAs were performed as previously described ^2^ using coating and biotinylated detection antibodies from R&D or Biolegend. Streptavidin-conjugated HRP was used for detection with a commercially available substrate solution, and the results were detected on a Molecular Devices SpectraMax Microplate Reader. Homogenates of lung tissues were prepared using a Polytron homogenizer and protein concentrations for all samples were equalized prior to ELISA.

### Statistics

The results below are expressed as either individual mice/data points or group means ± standard deviation (SD). P values and significances were determined using either the one-tailed Mann-Whitney T-test or non-parametric one-way ANOVA (GraphPad Prism© software). Groups were judged to be significantly different if the P value was less than 0.05 (*:p<0.05, **:p<0.01, ***:p<0.001).

## Notes

### Competing Interest Statement

The authors have declared no competing interest.

